# Molecular characterization of SARS-CoV-2 in the first COVID-19 cluster in France reveals an amino-acid deletion in nsp2 (Asp268Del)

**DOI:** 10.1101/2020.03.19.998179

**Authors:** Antonin Bal, Grégory Destras, Alexandre Gaymard, Maude Bouscambert-Duchamp, Martine Valette, Vanessa Escuret, Emilie Frobert, Geneviève Billaud, Sophie Trouillet-Assant, Valérie Cheynet, Karen Brengel-Pesce, Florence Morfin, Bruno Lina, Laurence Josset

## Abstract

We present the first genetic characterization of a COVID-19 cluster in Europe using metagenomic next-generation sequencing (mNGS). Despite low viral loads, the mNGS workflow used herein allowed to characterize the whole genome sequences of SARS-CoV2 isolated from an asymptomatic patient, in 2 clinical samples collected 1 day apart. Comparison of these sequences suggests viral evolution with development of quasispecies. In addition, the present workflow identified a new deletion in nsp2 (Asp268Del) which was found in all 3 samples originating from this cluster as well as in 37 other viruses collected in England and in Netherlands, suggesting the spread of this deletion in Europe. The impact of Asp268Del on SARS-CoV-2 transmission and pathogenicity, as well as on PCR performances and anti-viral strategy should be rapidly evaluated in further studies.

---

To the Editor,

In December 2019, a novel coronavirus emerged in China, causing outbreaks of unexplained pneumonia [1]. The virus was subsequently identified as a beta-coronavirus and named severe acute respiratory syndrome coronavirus 2 (SARS-CoV-2). SARS-CoV-2 is responsible of the coronavirus disease 2019 (COVID-19) pandemic which includes asymptomatic, upper, and lower respiratory tract infections. Among the first European cases of COVID-19, 6 were associated with a cluster of transmission in the French Alps in late January 2020 [2]. The index case of this cluster travelled from Singapore to France and went back to the United Kingdom (UK) where he was tested positive for SARS-CoV-2 on February 6^th^. Here, we aimed to investigate the French cases related to this cluster using metagenomic next-generation sequencing analysis (mNGS).

Of the 6 contact patients tested positive for SARS-CoV2, the 3 samples with highest viral loads (assessed by RT-PCR targeting the RdRp gene) were selected for mNGS [3]. One nasopharyngeal swab was collected from a patient with an upper respiratory tract infection on February 7^th^ (sample #1, Ct = 31.3). The other two samples were collected from the same asymptomatic patient on February 8^th^ (sample #2, nasopharyngeal swab, Ct = 31.1) and 9^th^ (sample #3, nasopharyngeal aspirate, Ct = 28.8).

A previously described mNGS protocol was used but for which DNase treatment was performed after nucleic acid extraction in order to increase sensitivity of RNA viruses’ detection [4]. Low quality and human reads were filtered out and remaining reads were aligned to the SARS-CoV-2 reference genome (isolate Wuhan-Hu-1, EPI_ISL_402125) using BWA-MEM algorithm. A mean of 19 445 767 reads per sample were generated, of which a mean of 605 243 reads per sample were mapped to the SARS-CoV-2 reference genome. Percentage of genome covered at a minimum depth of coverage of 100X was 38.3% for sample #1, 99.6% for sample #2 and 80.6% for sample #3.

The whole genome sequence (WGS) generated from sample #2 was deposited on GISAID (EPI_ISL_410486). The phylogenetic analysis using the 571 WGS of SARS-CoV-2 publicly available (as of March 17^th^ 2020) found that this sequence clustered with a sequence collected in Jiangsu, China (EPI_ISL_408488) on January 19^th^, suggesting a direct introduction of the virus from Asia, as suggested by epidemiological data (Fig 1). Compared to the reference SARS-CoV-2 sequence, a 3-nucleotide deletion in open reading frame 1ab (ORF1ab) at position 1607-1609 was identified. This deletion was found in 100% of the reads at this position with a coverage of 1745X around the deletion. Importantly, this deletion was also identified in 100% of the reads of sample #1 and sample #3 with a coverage of 54X and 481X, respectively. Using CoV-GLUE resource, we found that this mutation leads to a deletion of amino acid 268 in non-structural protein 2 (nsp2) [5]. Up to March 17^th^, this deletion in nsp2 (Asp268Del) was also characterized in 37/571 (6.1%) WGS (England n = 6; Netherlands n = 31). WGS-based phylogenetic analysis found that 15 viruses containing this specific deletion were closed to viruses collected in China between December 2019 and early February 2020, while 23 viruses collected in the Netherlands in March 2020 have slightly diverged (Fig 1).

**Figure 1.**
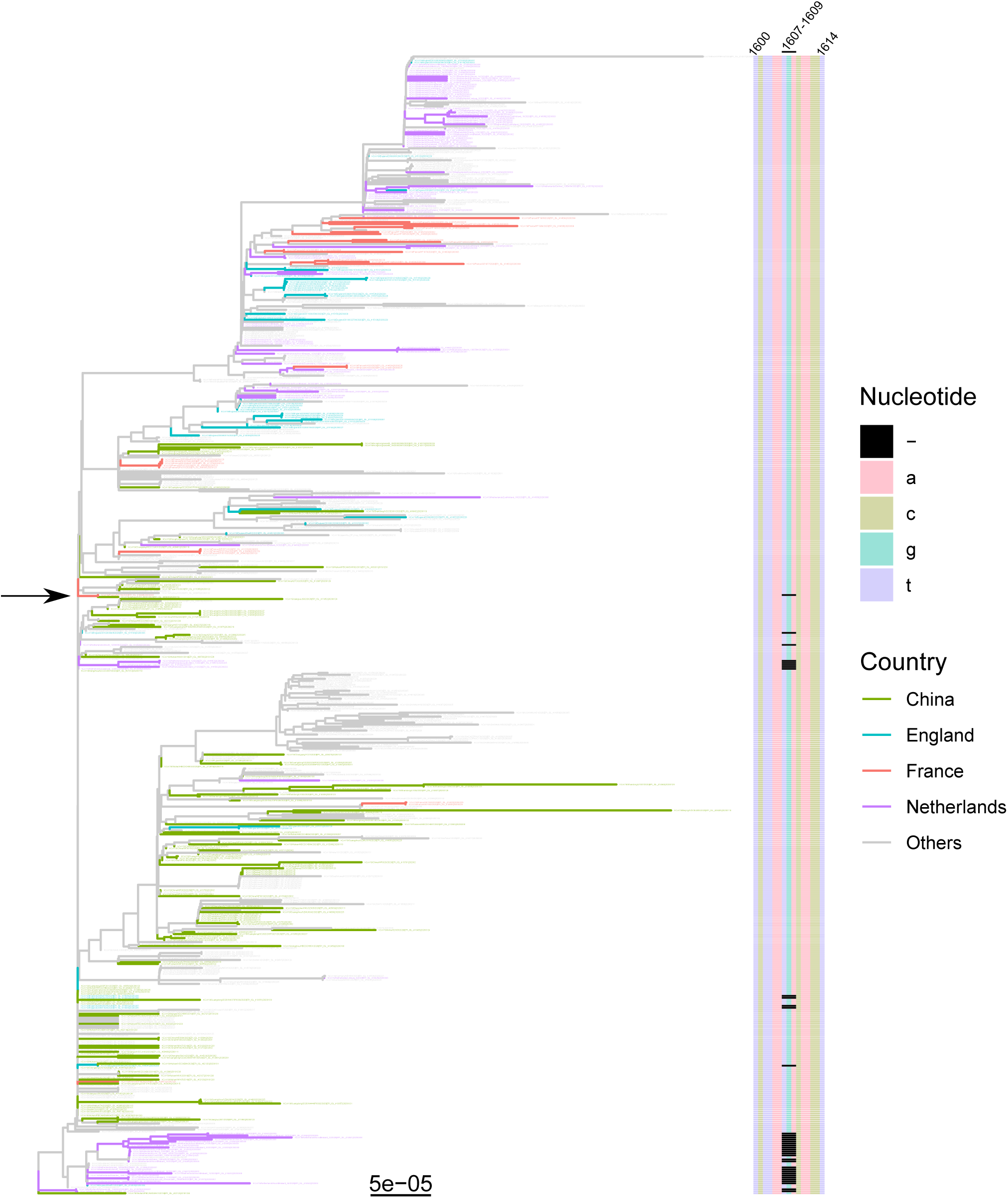
WGS-based phylogenetic analysis of SARS-CoV-2. The analysis included 571 WGS of SARS-CoV2 (>29k nt) collected in humans and available in GISAID from March 17^th^, 2020. The following sequences were excluded from the analysis: EPI_ISL_406592, EPI_ISL_414588, EPI_ISL_412900, EPI_ISL_408487, EPI_ISL_408483 and EPI_ISL_406595 because they were outliers, and EPI_ISL_413747, EPI_ISL_413695 because of incomplete sequences in ORF1ab. The hCoV19/Wuhan/IPBCAMSWH01/2019 strain was used as an outgroup virus. Genetic distances were calculated using the Kimura’s 2-parameter model (K80) and pairwise deletion. The tree was constructed by the neighbour-joining method using R seqinr and ggtree packages and validated using 1000 bootstrap pseudo-replicates. Sequence from sample #2 (EPI_ISL_410486) is shown by the arrow. Nucleotide alignment (1600 to 1614) is depicted as a heatmap on the right panel with 3-nucleotides deletion shown in black.

SARS-CoV2 sequences were not further compared between the 2 patients due to largely incomplete coverage of SARS-CoV2 genome in sample #1. Nonetheless, the longitudinal samples from the asymptomatic patient (sample #2 vs sample #3) were compared using a minimum depth of coverage of 100X in order to make a preliminary assessment of intra-host genetic variability. Three SNVs were noticed between the 2 samples: C366A (nsp1: S34Y), A20475G (synonymous mutation in nsp15), T24084A (protein S: L841H), suggesting intra-host evolution of the virus. For all 3 positions, nucleotides from sample #2 were still detected in sample #3 but as minor variants.

In this short report, we present the first genetic characterization of a COVID-19 cluster in Europe. Despite low viral loads, the mNGS workflow used herein allowed to characterize the whole genome sequences of SARS-CoV2 isolated from an asymptomatic patient, in 2 clinical samples collected 1 day apart. Comparison of these sequences suggests viral evolution with development of quasispecies. Specific studies using high depth of coverage to assess quasispecies are needed to characterize potential intra-host adaptation. In addition, the present workflow identified a new deletion in nsp2 (Asp268Del) which was found in all 3 samples originating from this cluster as well as in 37 other viruses collected in England (February) and in Netherlands (March), suggesting the spread of this deletion in Europe. The impact of Asp268Del on SARS-CoV-2 transmission and pathogenicity, as well as on PCR performances and anti-viral strategy should be rapidly evaluated in further studies.

## Ethical considerations

Investigations complied with the General Data Protection Regulation (Regulation (EU) 2016/679 and Directive 95/46/EC) and the French data protection law (Law n°78-17 on 06/01/1978 and Décret n°2019-536 on 29/05/2019). Informed consent to disclosure information relevant to this publication was obtained from the confirmed cases in France.

## Acknowledgements

We would like to thank all the patients, laboratory technicians, and clinicians who contributed to this investigation. We are also grateful to Véréna Landel and Philip Robinson Landel (DRCI, Hospices Civils de Lyon) for help in manuscript preparation. We thank the Authors, the Originating and Submitting Laboratories for their sequence and metadata shared through GISAID on which this research is based. We gratefully acknowledge all the members of CoV-GLUE, Nextrsain.org, and virological.org for sharing their analysis in real-time.

